# Endogenous controllability of closed-loop brain-machine interfaces for pain

**DOI:** 10.1101/369736

**Authors:** Suyi Zhang, Wako Yoshida, Hiroaki Mano, Takufumi Yanagisawa, Kazuhisa Shibata, Mitsuo Kawato, Ben Seymour

## Abstract

The ultimate aim of closed-loop brain-machine systems for pain is to directly titrate the ongoing level of an intervention to pain-related neural activity. However pain is highly susceptible to endogenous modulation, raising the possibility that active or passive changes in neural activity provoked by the operation of the system could enhance or interfere with the signals upon which it is based. We studied healthy subjects receiving intermittent pain stimuli in a real-time fMRI-based closed-loop feedback-stimulation task. We showed that multi-voxel pattern decoding of pain intensity could be used to train a control algorithm to learn to deliver less painful stimuli (adaptive decoded neurofeedback). However, the system engaged two types of endogenous processes in the brain. First, despite the inherent incentive for subjects to enhance the neural decodability of pain, decodability was either reduced or unchanged in classic pain-processing regions, including insula, dorsolateral prefrontal, and somatosensory cortices. However, increased decodability was observed in a putative pain modulatory region - the pregenual anterior cingulate cortex (pgACC). Second, we found that pain perception itself was modulated by an endogenous computational uncertainty signal engaged as subjects learned the success rate of the system in reducing pain - an effect that also correlated with pgACC responses. The results illustrate how regionally and computationally specific co-adaptive brain-machine learning influences the efficacy of closed-loop systems for pain, and shows that pgACC acts as a key hub in the endogenous controllability of pain.

## Introduction

The eventual goal of closed-looped brain-machine interfaces for pain is to use brain activity patterns to dynamically control a therapeutic intervention, such that the level of intervention is automatically tuned to the level of pain (Shirvalkar et al., 2018; Zhang and Seymour, 2014). Recent studies have showed that multivariate pattern analysis (MVPA ‘decoding’) methods can be used to discriminate pain intensity related brain (BOLD) responses in humans with reasonable accuracy (Brodersen et al., 2012; Marquand et al., 2010; Schulz et al., 2012; Wager et al., 2013), and so in principle pain-related brain activity can be used to guide interventions in closed-loop settings. However, if a person is aware that their brain is being decoded for this purpose, the incentive exists for them to try and endogenously enhance the representation of pain-related brain activity to more accurately signal to the machine decoder that the current intervention is working or not working. Whether this is achievable is not known, and it remains possible that attempts to enhance pain decodability could paradoxically interfere with it. Either way, it is clearly important for predicting the performance and stability of closed-loop systems over time.

The issue strikes at the heart of our understanding of the basic neural mechanisms of the endogenous control of pain. It is well recognised that the pain system is remarkably susceptible to subjective modulation by higher brain control processes (Basbaum and Fields, 1984). Neuroimaging studies have consistently shown that pain-related responses in a number of brain regions are increased or decreased in parallel with behavioural endogenous decreases (hypoalgesia) or increases (hyperalgesia) in pain (Atlas and Wager, 2012; deCharms et al., 2005; Tracey and Mantyh, 2007; Wiech, 2016; Woo et al., 2017). But this doesn’t necessarily imply that the discriminability of intensity on the basis of brain responses (i.e. neural decodability) is under control. If it is, it would indicate a much more specific dimension to endogenous modulation than previously known, because it would imply that the informational content (i.e the signal to noise ratio) of pain-related brain activity is under direct control, rather than a more non-specific up- or down-regulation of pain-related activity. It also raises the question as to what influence such attempts to modulate pain-related neural activity have on the subjective perception of pain.

To examine this issue, we designed an adaptive brain-machine interface system using real-time functional brain imaging (rtfMRI) linked to a machine that controlled an electrical pain stimulator. The aim of the machine was to learn to reduce the intensity of experienced pain based on decoded BOLD representations of pain in the brain. Specifically, the machine treated a low likelihood of decoded pain as its goal, so success would involve learning to select a lower current intensity to the stimulator. In this way, the machine acts in the best interests of the subject, as a protective system in which the level electrical stimulation is the intervention under closed-loop control. Importantly, subjects were made aware of the system architecture, creating the incentive for them to better communicate their pain (via their brain activity) to the machine: i.e. enhancing pain representations to better teach the machine when they are in pain (so that it would change the current intensity), or not in pain (so that it would not change the current intensity).

## Results

19 healthy participants completed the experiment, which took place over 2 days. On Day 1 (decoder construction sessions), participants received a sequence of high and low intensity painful electrical stimuli to the left hand, with no associated task demands other than intermittent pain ratings. For decoding, we used BOLD responses in bilateral insula cortex (incorporating posterior, mid and anterior insula), since this is thought to incorporate subregions that have a primary role in the coding of pain (Craig, 2002; Geuter et al., 2017; Segerdahl et al., 2015; Woo et al., 2017). Individual participant’s responses to high and low intensity stimuli were subsequently used to train a voxel-wise MVPA decoder that could classify the two stimulus levels.

On Day 2 (neurofeedback adaptive control sessions), high and low pain stimulation were embedded into a closed-loop adaptive control paradigm using real-time MVPA output as the feedback signal. Specifically, a machine controlling the two levels of pain stimulation tried to learn to deliver the level with the least pain based on the output of the MVPA classifier (the Day 2 scan was carefully realigned with Day 1 to allow decoder generalisation across days). The reason for using an adaptive decision function was two-fold: to maximise the incentive for subjects to enhance neural discriminability of pain (because the impact of successfully communicating the output is ‘remembered’ by the control algorithm for future trials); and to show that in principle, pain-reducing interventions can be *learned* by an appropriate control algorithm (i.e. the machine has no a priori knowledge that the lower current stimulus causes less pain, but is able to learn this from trial-and-error). Specifically, stimulation was based on a reinforcement learning algorithm that learned the values for each of the two stimulus levels, initialised at zero at the beginning of each session. After delivering a pain stimulus, it used the prediction from the MVPA classifier as the reward value in the RL algorithm (with lower probability of high pain equating to a more positive reward). Therefore, given any above chance performance of the MVPA classifier, the machine should learn to deliver low pain intensity stimuli more frequently.

To allow meaningful comparisons to be made between the brain responses on the two days, bearing in mind their necessary sequential order, we yoked a previous participant’s Day 2 pain sequence as the sequence for a future participant on Day 1. Furthermore, Day 1 and 2 had the same trial structure (Fig 1c): high or low intensity pain were delivered at the beginning of each trial, coinciding with a ‘+’ symbol appearing on the screen below the fixation point which remained for 10s, followed by a brief inter-trial interval (ITI) of 2s showing an ‘=’ symbol (Fig 1d). On 40% of all trials in (12 of 30 each session), the fixation point turned into an orange square, notifying the participants that their pain ratings would be taken soon. A visual analogue scale with 0 to 10 appeared 4s after trial start, and stayed on for 6s for pain rating. Brain images from 4-10s after pain delivery were used for both decoder training and real-time decoding. This allowed for the BOLD delay and avoided movement contamination.

**Figure 1:**
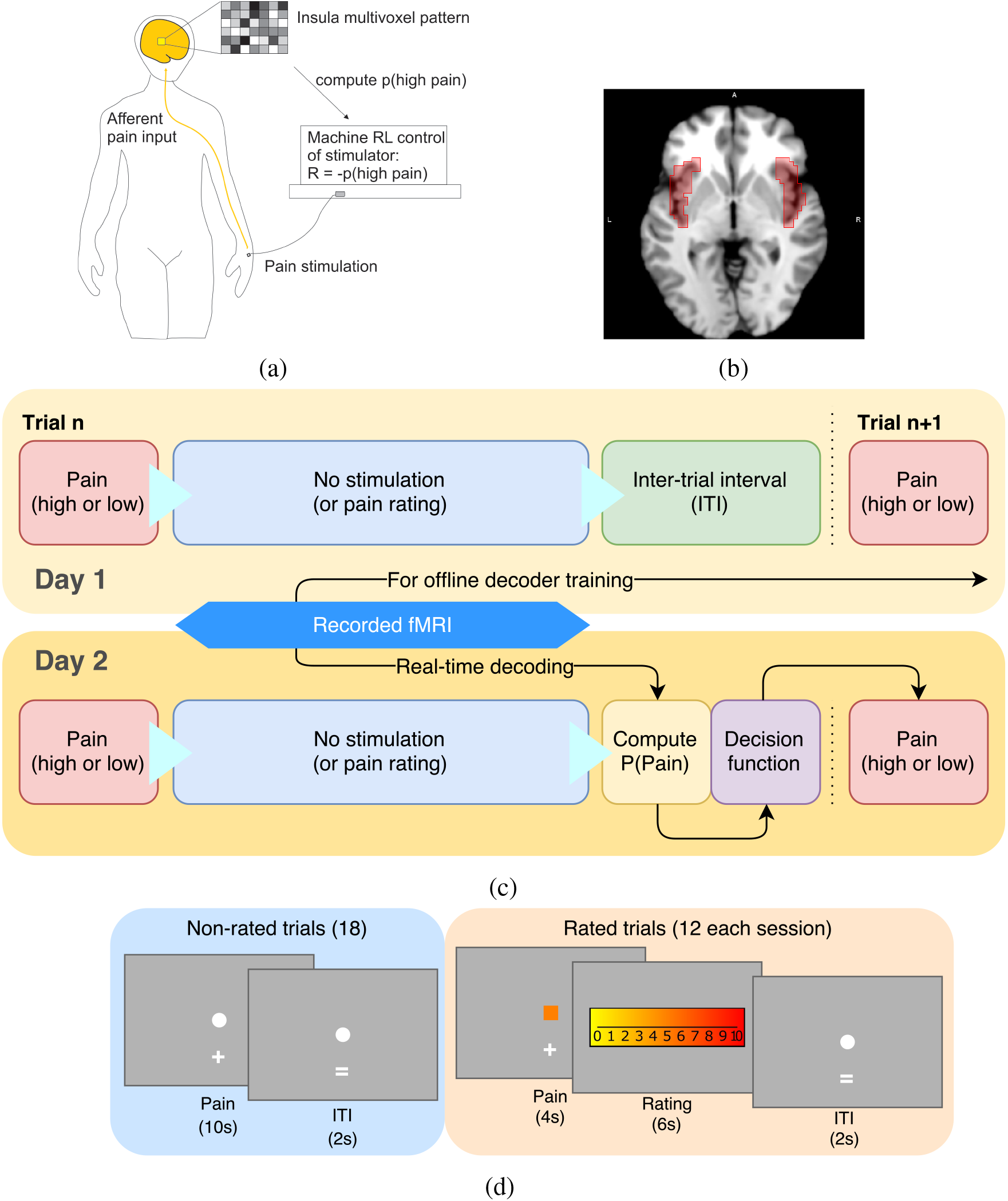
Experimental paradigm. (a) Schematic illustration of the closed-loop setting. (b) Demonstration of bilateral insula ROI generated from AAL atlas (viewed at z=0). (c) Participants took part in a two-day decoding neurofeedback experiment. On Day 1, their functional brain images were recorded while they were given high or low level of electrical pain at the beginning of each trial, which were used for offline MVPA decoder training. On Day 2, participant’s probability of having received high pain (P(Pain)) in the current trial was computed in real-time from their brain activities by the decoder, which was then used by the pain delivery system to decide on the pain level of the next trial. The decision function of the system was based on a reinforcement learning algorithm that aimed to lower the participant’s decoded P(Pain). (d) For 40% of all trials in a session, participants were asked for their pain ratings 4s after its delivery on a 0-10 VAS scale. The changed fixation point acted as a prompt for rating. The display were identical on both days.

On Day 1, subjects were informed that they would receive a random sequence of pain stimuli, and that their only task was to perform the intermittent ratings. On Day 2, subjects were fully informed about the closed-loop operation, and how the clarity of their brain responses would help the machine give them less pain (see instructions in Appendix). The goal was to illustrate the opportunity for subjects to modulate their brain activity, without providing any explicit instruction of what to do.

### Behavioural results

Within-subject decoder construction based on the insula ROI achieved reasonable classification accuracy (Day 1: 10-fold cross-validated test accuracy 65%, sensitivity 60%, specificity 67%, accuracy one-sample t-test vs 0.5 across subjects: T(18)=8.967, p<1e-7) (see also Tables 1). When this classifier was used during neurofeedback when the subject returned on Day 2, decoding accuracy remained above chance (Day 2: accuracy 56%, sensitivity 51%, specificity 63%, accuracy t-test vs 0.5: T(18)=4.053, p=0.0007). Specifically, the real-time decoder output following delivery of high pain (referred to as P(pain), Fig 2a) differed significantly for the high and low pain stimuli (repeated measure ANOVA of session and pain level effects, only pain level main effect significant: F(1, 18)=17.41, p=0.0006, post-hoc test Bonferroni corrected P(pain) for high pain 95%CI=[0.558, 0.676], low pain=[0.434, 0.551]).

**Table 1:** Decoder testing performance (high pain = positive, low pain = negative for sensitivity/specificity calculation; CV: 10-fold cross validation; D1: Day 1; D2: Day 2. All values are mean (SEM), n=19)

|  | Train D1, Test D1 (CV) | Train D1, Test D2 | Train D2, Test D2 (CV) | Train D2, Test D1 |
| --- | --- | --- | --- | --- |
| Accuracy | 0.6488 (0.0163) | 0.5632 (0.0156) | 0.5601 (0.0100) | 0.4912 (0.0314) |
| Sensitivity | 0.6017 (0.0258) | 0.5057 (0.0159) | 0.4982 (0.0313) | 0.4379 (0.0258) |
| Specificity | 0.6654 (0.0248) | 0.6309 (0.0365) | 0.5903 (0.0246) | 0.5490 (0.0310) |
| # features (voxels) | 24.053 (1.053) |  | 28.737 (0.700) |  |

**Figure 2:**
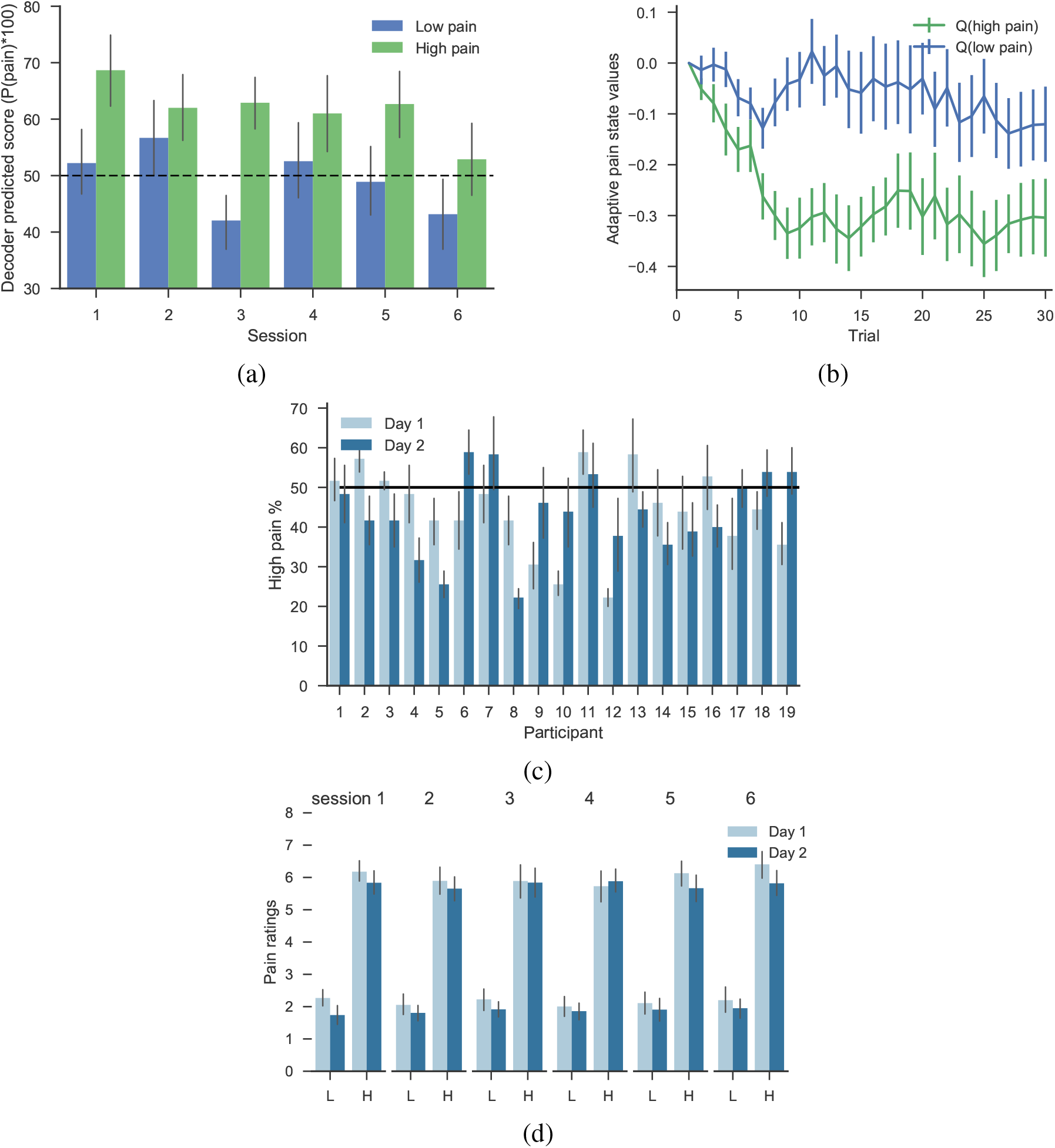
Behavioural results (mean ± SEM across n=19 individuals). (a) Decoder predicted probabilities of having received high pain, P(pain), were able to distinguish high/low pain state (calculated on Day 2 only). (b) Within-session, the control system learned to value low pain states higher than high pain states (Q(low pain) > Q(high pain)) after several initial trials (Day 2). (c) High pain trials were delivered less frequently than low pain (participant 1-3 used random stimulus sequences on Day 1 instead of yoked, SEM calculated from sessions). (d) Raw pain ratings were significantly different for the two pain levels, but not across sessions or days (H=high pain, L=low pain).

Decoder performance was therefore sufficient for the reinforcement learning control system to learn differential values for high and low pain stimuli within a few trials in each session (Fig 2b, mean±SEM, high pain=-0.264±0.0486, low pain=-0.0608±0.0479, paired t-test: T(18)=-3.651, p=0.0018). Accordingly, the control system delivered significantly fewer high compared to low pain stimuli (Fig 2c, high pain percentage: Day 1=44.123±2.394%, Day 2=43.480±2.353%, one-sample t-test vs 50%: Day 1 T(18)=-2.455, p=0.0245, Day 2 T(18)=-2.771, p=0.0126).

An important feature of our experimental design was to yoke sequences on Day 2 (neurofeedback) to Day 1 (decoder construction), to allow meaningful comparisons to be made. To achieve reasonable classification learning, therefore, we set the decision function (i.e. by which action values determine the actual machine choice of pain stimulation level) for the control system to be noisy, so that despite the relatively clear difference between action values, there were sufficiently large number of high stimuli delivered that would support decoder training performance when the sequence was used for a subsequent subject’s Day 1 decoder construction. Note that the 3 initial subjects did not have yoked stimulus sequences, instead they used randomly generated 50/50 high/low pain sequences.

Participants completed pain threshold testing on both days before the experiment started, with aims to achieve VAS=1 and 8 for low/high pain respectively. Consequently, pain ratings were significantly different between the high and low levels of pain (Fig 2d, high pain=5.92±0.356, low pain=1.99±0.258, repeated measure ANOVA Pain levels: F(1,11)=86.00, p<1e-5), but did not show significant differences across sessions or days (day: F(1,11)=3.173, p=0.103, session: F(5,55)=0.470, p=0.797). There were no significant differences in the high/low pain stimulation current levels given between days (paired t-test p=0.12 and 0.27 respectively).

We conducted a questionnaire survey after Day 2’s experiment concluded, in which participants showed evidence of awareness of and attempt in mentally influencing the system to reduce overall pain. 17 out of 19 (1 ambiguous) of all participants believed the machine was successful in reading their pain and trying to reduce it, 15 out of 19 (2 ambiguous) believed that they were successful in influencing the system to achieve that using some mental strategies. The strategies used most frequently included a combination of mental imagery of pain, distraction from pain, predicting/recalling stimulus sequence, and doing nothing.

### Whole-brain BOLD comparison between days

Offline whole-brain analysis of fMRI data using a conventional general linear model showed evidence of a regional day × pain level interaction (Fig 3a). Specifically, within-subject comparison (Day 2 > Day 1) of the contrast (high pain > low pain) showed increased responses in periaqueductal grey (PAG) (statistics in figure legend, and correction for multiple comparisons detailed in Tables 2), in a region most likely localising to dorsolateral or lateral PAG (Fig 3b), using PAG subdivision masks from (Ezra et al., 2015). The PAG is an important *a priori* region of interest in this analysis, as it is a key part of the descending endogenous control system. We did not observe responses elsewhere at whole-brain corrected thresholds.

**Figure 3:**
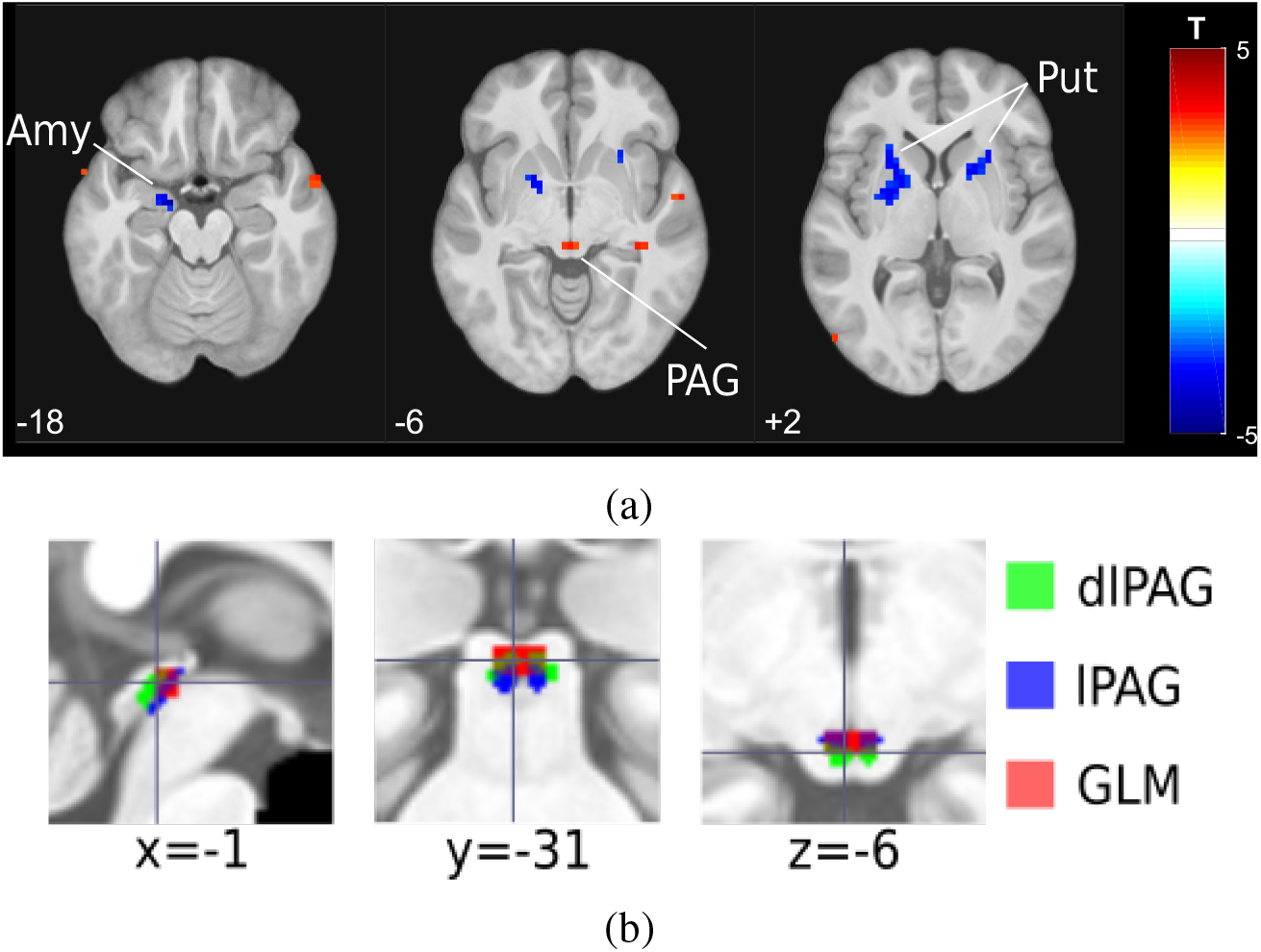
Whole brain comparison between days results (n=19), (a) Within-subject comparison (2nd level paired t-test, Day 2 > Day 1) of the high pain > low pain 1st level contrasts, interaction were observed in left amygdala (peak coordinates [−16, −7, −18], T=-4.38, k=11), bilateral putamen (left peak [−26, 12, 6], T=-4.09, k=47, right peak [26, 19, −2], T=-4.50, k=32) and the PAG (peak coordinates [0, −30, −6], T=3.27, k=3). All images shown at p<0.005, k>0. (b) Dorsal lateral PAG and lateral PAG masks (thresholded at 50%) from (Ezra et al., 2015) overlaying PAG activation from GLM (cluster formation at Z=2.9). SVC using bilateral dlPAG mask p(FWE-corr)=0.034, k=1, T=3.14, Z=2.76, peak in [−3,-30,-6], k=1, T=2.94, Z=2.62, [3,-30,-6], lPAG mask (FWE-corr)=0.036, other statistics the same as dlPAG mask. (note our voxel size is too big for very serious PAG subdivision). H: high pain, L: low pain.

In contrast, we found decreased BOLD responses in the left amygdala and bilateral putamen (Fig 3a). Responses in the bilateral insula ROI, of which the decoder computed P(Pain) from, were not significantly different between days for either high or low pain, or overall (pain level main effect: F(1,18)=63.911, p=2.475e-7, session main effect: F(5,90)=4.130, p=0.002, none of the interactions significant. Note that images used in the GLM and subsequent analyses were fully preprocessed, as opposed to the limited (i.e. rapid) processing used in real-time neurofeedback.

### Decoder comparison

To identify potential changes in the multi-voxel patterns in the insula, we compared MVPA decoder performance trained offline using the bilateral insula ROI on functional brain images from Day 1 and Day 2 (i.e. training the decoder separately on each day, and using internal cross-validation to test performance). This analysis aimed to detect changes in the representation of pain, distinct from the mean BOLD signal across all ROI voxels, and despite the fact that there were no significant changes to pain stimuli or rating between days. Using a 10-fold cross-validation, we found that the decoding test accuracy decreased on day 2 compared to Day 1 (Fig 4a and Tables 1, Day 1: 64.8%, Day 2: 56.0%, Wilcoxon signed-rank test Z(18)=3.69, p<0.001). Furthermore, the decoder trained with Day 2 data identified significantly more voxels as contributing to decoding performance compared to Day 1 (Fig 4b and Tables 1, Day 1: 24.1±1.1 voxels, Day 2: 28.7±0.7 voxels, signed-rank test Z(18)=-3.21, p=0.001) (NB. the sparse logistic regression method we used prunes unnecessary features when building the classifier Yamashita et al. (2008)). There were no significant differences in the locations (i.e. average of x, y, or z coordinates) of these weighted voxels within individuals. These results suggest the insula as an ROI may have overall disrupted functional information content for pain level encoding on Day 2 (neurofeedback).

**Figure 4:**
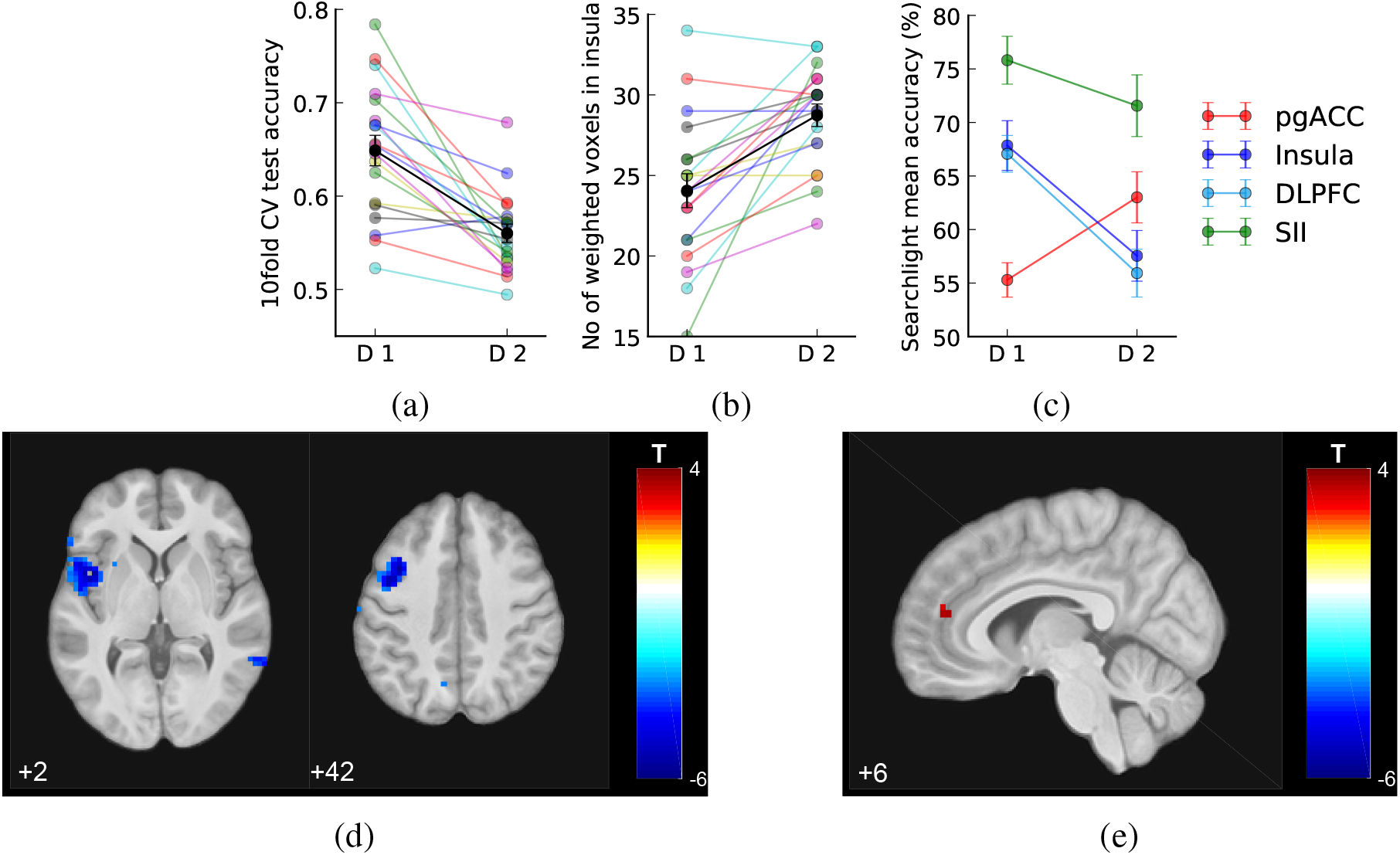
Decoder comparison and searchlight analysis results (mean SEM across N=19 individuals). (a) 10 fold cross validation test accuracy of insula decoder decreased on Day 2. (b) Number of weighted voxels increased on Day 2. (c) Comparison between the mean searchlight accuracy of pgACC and insula clusters (masks extracted from clusters shown in figures below, at p<0.005 unc.) and right SII (mask from accuracy>75% on Day 1, k=50 voxels, see text). (d-e) Whole-brain searchlight analysis showed that information content contributing to decoding accuracy decreased in left insula and DLPFC/MFG, and increased in pgACC, on Day 2 comparing to Day 1 (shown at p<0.005, k>0).

Therefore to look more broadly across the whole brain and identify any changes in pain intensity representations, we conducted a whole brain post-hoc searchlight analysis using data from Day 1 and 2. This identifies accuracy maps that reflect the local information content of each voxel (Hebart et al., 2015; Kriegeskorte et al., 2006), which can be used to search the brain for changes in pain level representation. A paired t-test of these maps (Day 2 > Day 1 in a second level paired t-test, DF=18) revealed reduced pain level decoding accuracy localised to a region in the left mid/anterior insula (Fig 4d, Tables 2, [−45, 6, 2], T=-6.04, k=142, whole brain cluster level p(FWE-corr)=0.014, shown at p<0.005 uncorrected). This localisation presumably underlies the ROI-based reduction in accuracy in the preceding analysis. Extracting the exact values from accuracy maps from both days, the left insula showed decreased decoding accuracy on Day 2 (171 voxels, Day 1: 67.844±2.320, Day 2: 57.546±2.366, paired t-test T(18)=-5.335, p=4.525e-5). Outside of the insula, we also noted reduced accuracy (135 voxels, Day 1: 67.074±1.715, Day 2: 55.932±2.234, paired t-test T(18)=-4.996, p=9.359e-5) in the left medial frontal gyrus (i.e dorsolateral prefrontal cortex (DLPFC), [−38, 9, 42], T=-5.68, k=134), which survived correction for whole brain multiple comparisons (whole brain cluster level p(FWE-corr)=0.045).

In contrast, using the same searchlight method as above, we found *increased* information content in a small region consistent with the pregenual anterior cingulate cortex (pgACC) in the medial prefrontal cortex (Fig 4e, Tables 2, [6, 44, 14], T=3.50, k=5, small volume correction (SVC) using an 8-mm spherical mask based on our previous investigation (Zhang et al., 2018)). Extracting the exact values from the accuracy maps from both days, this pgACC ROI had significantly increased decoding accuracy across all participants (Fig 4c, Day 1 accuracy: 55.293±1.604, Day 2: 63.009±2.383, paired t-test T(18)=3.676, p=0.0017). No other brain regions were identified, even at a low exploratory threshold.

For comparison, we also looked at the average accuracy maps from all participants in the right secondary somatosensory cortex (SII), which had the highest decoding accuracy on both days. (Day 1 peak [45,-17,26], accuracy=77.414, Day 2 peak [55,-30,26], accuracy=74.510). We found that SII decoding accuracy did not vary significantly across days (averaged within the cluster mask from Day 1, k=50, Day 1: 75.813±2.234, Day 2: 71.563±2.880, paired t-test T(18)=1.344, p=0.196). To look at regional differences formally, we computed a day × location interaction: we found that this was significant between pgACC and SII (F(1,18)=6.648, p=0.012), although not significantly so between insula and SII (F(1,18)=1.507, p=0.22).

### ‘Switch’ trials analysis

The imaging results above indicate that there are differences in the way pain is processed and represented on Day 2. To examine this further, we next looked for behavioural and brain evidence that might reflect the engagement of active (given the self-reported attempts) or passive modulation of brain activity in the context of the closed-loop neurofeedback. Specifically, we looked at ‘switch’ trials, in which the pain level delivered on the current trial differed from that of the previous trial (i.e. as the machine switched from high to low pain, or vice versa), because these trials carry information relevant to the success of the machine, and any mental strategy the subjects might use to try and influence it.

Using a simple t-contrast, we found that pain ratings for the low pain stimulus on switch trials was significantly higher than non-switch trials on Day 2, although there was no significant interaction across days (Fig. 5a, paired t-test T(18)=-2.466, p=0.0186, day × switch interaction for low pain: F(1,18)=2.477, p=0.133, interaction for high pain: F(1,18)=0.214, p=0.650). Following the same pattern, we found that Day 2 predicted insula P(Pain) score on switch trials was significantly higher for low pain (Fig 5b, paired t-test T(18)=2.990, p=0.0079), as well as being marginally so for high pain (T(18)=1.952, p=0.067). This provides evidence, from both ratings and multi-voxel insula patterns, that subjects may be sensitive to the sequence of pain stimuli on Day 2.

**Figure 5:**
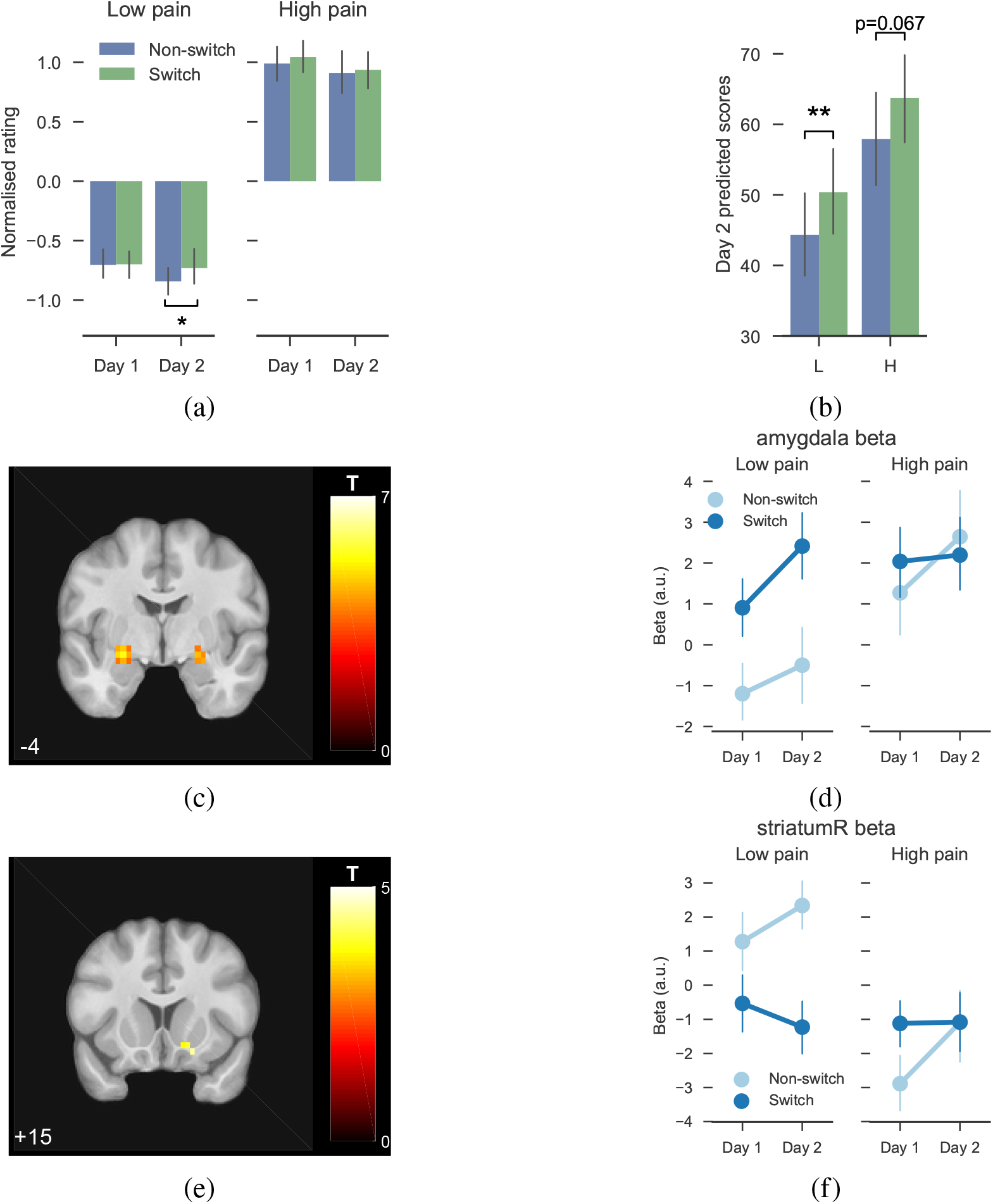
Switch trials differences (n=19). All imaging results shown at p<0.001, k=0. (a) Within-subject normalised ratings, group by days, pain levels, and switch status, showing that Day 2 switch low pain trials were more painful than non-switch trials. (b) Day 2 decoder predicted scores (p(pain)*100) for switch/non-switch trials showed differences for high and low pain. (c) Day 2 HL>LL in bilateral amygdala. (d) Beta values extracted using bilateral activation cluster as ROI (at p<0.001 unc., k=30). (e) Day 2 LL>HL in right ventral striatum / OFC. (f) Beta values extracted from individuals using striatum activation cluster (k=9).

In a basis GLM analysis of imaging data, the contrast of switch vs non-switch low pain trials on Day 2 revealed BOLD contrasts for HL>LL (i.e. a low pain stimulus trial that was preceded by a high stimulus, versus one preceded by another low stimulus) in bilateral amygdala (Fig 5c, Tables 2), and LL>HL in right striatum (Fig 5e). Exploring this result with an ROI analysis, Day 2 showed significant pain level × switch interaction in both ventral striatum (Fig 5f, repeated measure ANOVA F(1,18)=7.673, p=0.0126)), and amygdala (Fig 5d, ANOVA F(1,18)=11.991, p=0.0028)). This indicates that brain regions commonly associated with feedback learning appear to track pain feedback based on the machine-determined stimuli on Day 2.

### Frequency learning analysis

These basic switch trial analyses provide behavioural and brain evidence that subjects are sensitive to the sequential identity of the stimuli on Day 2 (although this effect is not readily apparent on Day 1, the lack of an interaction by day in the above analyses means we can’t necessarily conclude that subjects are *significantly* more sensitive to switches on Day 2). Switch trials are important because they contain more information than non-switch trials, an effect that can be formalised by a simple model in which people use the underlying frequency of high and low pain to infer how successful, overall, the machine is at reducing pain, i.e the overall probability of receiving low or high pain on any trial.

To capture a basic frequency learning process we applied a simple Bayesian learning model to quantify two key metrics: the ongoing probability of low/high pain, and the level of surprise on each trial (entropy). Previous studies have shown that such simple models provide a good account of behavioural and brain measures of surprise in comparable statistical learning environments (Mars et al., 2008; Meyniel et al., 2016).

We first looked at whether these metrics correlated with behaviour. Using a linear regression model of pain ratings (see methods), we found no correlation with *a posteriori* probability of low pain (z-transformed correlation coefficients Day 2 vs 0: T(18)=-0.582, p=0.568, Day 1 vs 0: T(18)=0.233, p=0.819, paired t-test between days: T(18)=0.601, p=0.555). However, we found a strong correlation with entropy, which was specific to Day 2, compared to Day 1. That is, greater entropy was associated with greater subjective pain (Fig 6a, z-transformed correlation coefficients Day 2 vs 0: T(18)=4.648, p=1.99e-4, Day 1 vs 0: T(18)=0.259, p=0.798, Paired t-test between days: T(18)=2.245, p=0.0376).

**Figure 6:**
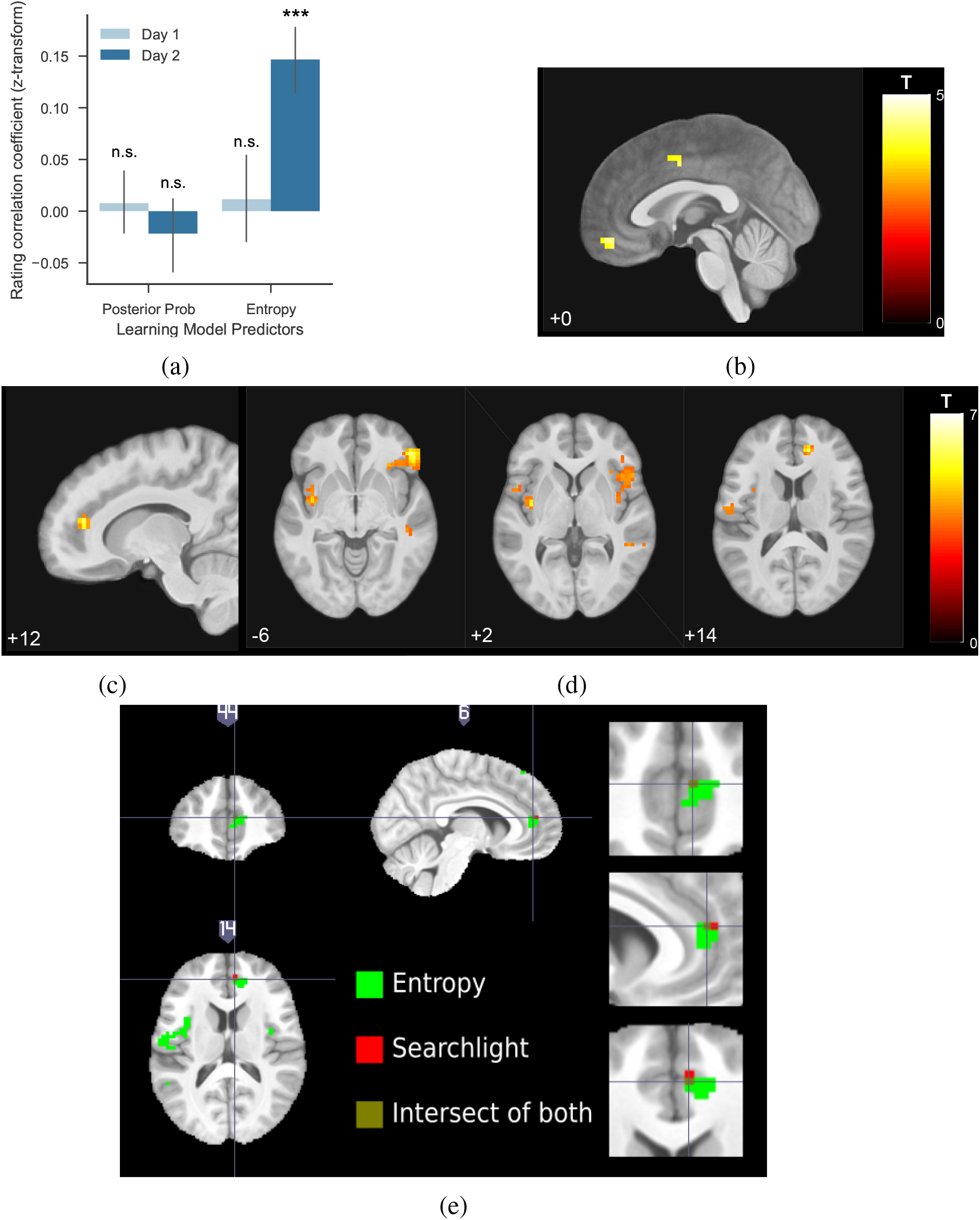
Frequency learning evidence on Day 2 (n=19). (a) Entropy (uncertainty regarding upcoming stimulus being high pain) from frequency learning model correlated with pain rating residuals from Day 2, but not Day 1 (using pain rating residuals with intensity and session numbers regressed out, see Methods). (b) Frequency learning model posterior probability of low pain correlated with VMPFC on Day 2 (peak coordinates [0, 51, −14], T=4.44, shown at p<0.001 unc.). (c-d) Frequency learning model entropy on Day 2 (i.e. entropy of posterior probability of current stimulus before updating) correlated with pgACC and bilateral insula (pgACC peak coordinates [13, 41, 14], T=5.91, sagittal and coronal views both at p<0.001 unc.). (e) Overlay of pgACC activations from both entropy (green) and searchlight (red) analysis (visualised at Z>3.2 for both).

In the analysis of imaging data on Day 2, we found that the *a posteriori* probability of low pain was correlated with BOLD responses in the ventromedial prefrontal cortex (VMPFC) (Fig 6b, Tables 2), consistent with this regions strong association with reward and relief value in previous studies of learning (Kim et al., 2006; Seymour et al., 2012). Most importantly (given the correlation with pain ratings), we found that entropy was correlated with BOLD responses in both bilateral mid/anterior insula and pgACC (Fig 6c and 6d, Tables 2). The insula response lay within the bilateral insula mask used for the decoder construction, and the pgACC response was part overlapping with the region associated with increased accuracy coding during adaptive neurofeedback (Fig 6e). When looking at the contrast of these responses across days, we found that although there was no significant effect of day in the insula response. However, the peak pgACC response was significantly greater on Day 2 (SVC corrected p(FWE-corr)=0.021, T=3.70, Z=3.15, peak coordinates [13,41,14]). That is, entropy correlated with both pain ratings and pgACC BOLD response selectively during neurofeedback on Day 2.

## Discussion

The results show that regional brain responses decoded in real-time can be used as a feedback signal to teach a machine to reduce the intensity of experimental pain stimuli in an adaptive, closed-loop setting. However, this is accompanied by two types of endogenous processes that influence both the neural signals themselves, and the perception of pain. First, in the context of the explicit incentive to enhance neural decodability, some primary pain-related brain regions (i.e. the insula) show a paradoxical decline in decodability, and the only region identified as showing enhanced decodability was the pgACC. Second, subjects learn the performance of the machine in terms of the success giving low intensity stimuli, and an entropy (uncertainty) signal evoked during learning is positively correlated with pain - an effect that was also associated with pgACC responses. The results illustrate how different, specific co-adaptive brain processes are engaged during the operation of closed-loop systems for pain.

In the context of closed-loop systems, the different patterns of adaptive response illustrate that different brain regions will generate feedback signals that may affect the performance of closed-loop systems in different ways. In the case of the experimental electrical stimulus here, secondary somato-sensory cortex performed best - with the highest decoding accuracy and resistant to the degradation of decodability seen in insula. It is not clear why insula shows a reduced performance, but previous evidence suggests that uncertainty (which correlates with a sub-regions of insula in our data) is integrated into perceptual representations as pain becomes more predictable (Brown et al., 2008; Geuter et al., 2017), and it may be that such a predictive coding schema disrupts decoder performance. It is notable, however, that overall the insula still predicts the slight increase in pain associated with changes in pain feedback (i.e. the switch trials, Fig 5b), so the results are still consistent with a primary role of insula in the subjective perceptual representation of pain.

However, the pattern of changes in the insula contrasts with that of the pgACC. Overall, the decoding accuracy in pgACC is much lower, consistent with the fact that it is considered not to be a primary pain coding region, but a potential center for endogenous control. From an ecological perspective, endogenous pain control is thought to provide a key mechanism by which animals cope with threat, and is mediated in part by control of descending pathways to the spinal dorsal horn neurons (via the PAG) that transmit incoming nociceptive signals (Basbaum and Fields, 1984). Along with other regions (including the insula and DLPFC), the pgACC has been implicated in endogenous control across a range of paradigms, including placebo/nocebo (Bingel et al., 2006; Eippert et al., 2009; Zubieta et al., 2005), attention/distraction (Bantick et al., 2002; Tracey et al., 2002; Valet et al., 2004), and controllability (Salomons et al., 2007; Zhang et al., 2018). But functionally dissociating the role of these different regions in endogenous modulation has been difficult, coupled with the fact that it has remained unclear whether modulation reflects a non-specific up- or down-regulation of pain responses, or as we now show here, a process in which the specific informational representation of pain is under control.

Specifically, the data here show that pgACC correlates both with a pain-modulating effect of uncertainty (entropy), and an increase in decodability under an appropriate explicit incentive. Importantly, intensity coding is needed for the computation of entropy, and thus the enhancement in decodability may reflect the enhancement of information used for this computation, ultimately leading to an influence on the subjective perception of pain. This is consistent with an attention-like process engaged specifically during neurofeedback, to enhance pain information processing in the context of the task. This computationally precise modulation is distinct from the basic representation of pain itself, and potentially mediated via the descending system given the non-specific enhancement of PAG activity seen during neurofeedback. Overall, the results show that pgACC may act as a key hub for neural and behavioural components of the endogenous control of pain, modulating the level of pain according to the informational value it carries in terms of its ability to guide active learning and behaviour.

From a neuro-engineering perspective, the experiment demonstrates that in principle, online decoded pain responses can guide a closed-loop pain control system, and the use of fMRI allows exploration of a range of target regions for decoding. Although multi-ROI classifiers can have much higher accuracy (Wager et al., 2013), here we arbitrarily used a single (bilateral) brain region, partly as it is more realistic in terms of future applications that involve long-term implanted recordings, for example with electrocorticography (ECoG). However, the choice of region is somewhat irrelevant to the experimental demonstration of endogenous processes studied here, as all that matters is that the decoder performs above chance and the incentive for active endogenous modulation exists.

As a general finding, it is clear that the brain actively learns about feedback in the context of brain-machine systems. Our experimental design here is relatively simple, and frequency learning is sufficient to capture the efficacy of the system. However, the engagement on brain regions including the striatum and amygdala suggests that more sophisticated value learning might be possible if the machine policy were more complex (for example, complex markov state-transition probabilities (Wang et al., 2017)). It is also possible that brain representations might change over extended periods of time based on the reinforcement provided by the feedback - and such ‘neural conditioning’ has been observed over multi-day decoded neurofeedback tasks that involve explicit reward feedback (Koizumi et al., 2016). It is therefore possible that these additional types of learning could further influence closed-loop systems in appropriate situations.

A key feature of our system is the incorporation of an reinforcement learning decision function on the part of the machine. This has a key advantage over fixed feedback decision policies because in principle RL algorithms can be used to search a much larger parameter space, as opposed to the binary levels of stimulation here - something that has broad applicability for many therapeutic interventions (e.g. spinal or brain stimulators). That is, when the optimal configuration of parameters for treatment under control is not known, the RL algorithm can search and find it over time. Combining the use of machine learning to generate a value approximation function, with reinforcement learning for optimal control, provides an ‘intelligent’ control systems approach to pain therapeutics.

## Methods and Materials

### Participants

19 healthy participants enrolled in a two-day neuroimaging experiment (two female, age 23.5±4.0 years). All subjects gave informed consent prior to participation, had normal or corrected to normal vision, and were free of pain conditions or pain medications. The experiment was approved by the Ethics and Safety committee of the Advanced Telecommunications Research Institute, Japan.

### Experimental protocol

The experiment spanned two days. On each day, each participant completed 2 sessions of pain thresholding test outside the scanner and 6 sessions of task with high/low painful stimuli inside the scanner.

#### Day 1: Decoder construction

Individual participant’s functional brain images were recorded during fMRI scanning for decoder training. High and low levels of painful electrical stimuli, determined with the participant’s pain threshold obtained before task outside the scanner, were delivered in a sequence of random or pseudo-random trials to elicit two levels of pain. From the participant’s perspective, painful stimulus was delivered at the beginning of each trial when a ‘+’ symbol appear on screen below the white bulls-eye fixation point. The ‘+’ stayed on for 10s, then the ‘=’ symbol replaced it for 2s, signalling a brief inter-trial interval (ITI). In 40% trials (12 randomly chosen out of 30 in each session), the ‘+’ stayed on screen for 4s and the fixation point turned to an orange square (signalling upcoming rating), followed by a 0-10 visual analogue scale that stayed on for 6s, where participants were asked to rate how painful the stimulus was by pressing two buttons to move the slider on screen. The 30-trial session was repeated 6 times with a short break in between (180 trials, 72 ratings per subject in total).

16 out of 19 participants used another participant’s Day 2 trial sequences on Day 1 as yoked control. All participants were given the instruction to rest in the scanner and do nothing (see ‘Appendix’).

Individual-specific, multi-voxel decoder was then trained for automatic classification of pain level experienced, using bilateral insula as region of interest (ROI, see ‘Decoder construction’).

#### Day 2: Neurofeedback adaptive control

On Day 2, the level of pain stimuli delivered on each trial was controlled by their decoded pain from real-time brain activities in the previous trial, determined by an adaptive control algorithm. All participants were explicitly told that the pain level they received was controlled by the computer programme, and were aware that modulating their brain activity could therefore influence the computer. The instructions are detailed in the Appendix, and were intended to reveal the incentive to modulate pain, but without any explicit instruction whether or how to do so.

After delivering pain, the participants’ probability of experiencing high pain (P(Pain)) was estimated by multiplying decoder weights with insula BOLD activity from their brain images in that trial (realigned and resliced to the reference image from Day 1, following Shibata et al. (2011)). The estimated probability was used to provide the feedback signal with the aim that the computer could learn to lower the overall level of pain delivered to the participant, based on trial-by-trial updating of the values of high and low pain stimulus with a basic reinforcement learning algorithm. The details of this algorithm are described below, but in brief, the stimulation state that elicited lower decoded pain signal in the participant was reinforced (see ‘Neurofeedback adaptive control’).

Day 1 and 2 were structurally the same apart from the adaptive control process and subject instructions, which made them approximately yoked conditions that allowed investigation of whether any brain-machine co-adaptation processes took place. Across any analysis of effect × day interactions, this sequential comparison necessarily introduces an order confound related to possible non-specific effects of novelty and anxiety to the experiment. In these instances, they are partly mitigated by the computational specificity of the analyses, and the fact that the majority of effects of interest emerge on day, during neurofeedback, when novelty and anxiety effects would be less.

### Stimulus delivery

Painful electrical stimuli were delivered using two constant current stimulators (Digitimer model DS7A, Welwyn Garden City, Hertfordshire, UK), at two current levels for high/low pain determined using the participant’s own threshold. The levels were fixed across sessions (except in 4 subjects, minor adjustments were made where pain ratings were either too high, or there were no difference between two levels), but were allowed to differ on Day 2 based on the new pain threshold. All stimuli were delivered as a train of 50 5ms square wave pulses at 10Hz, lasting 500ms (DS7 settings: x1 mA, 200*μ*s).

The two stimulators were connected to a custom-made switch that allowed current delivery through the same custom-made, MRI-compatible ring electrode (10mm diameter). The electrode was taped to the back of the left hand of the participant, its location marked on Day 1 as reference for attachment on Day 2.

### Pain thresholding procedure (Day 1 and 2)

On each day, participants completed a thresholding procedure at the beginning of the experiment. In the first session, the staircase method was used to evaluate their highest pain limit. Stimuli current were linearly increased at 0.2-0.5mA interval, and participants were asked for verbal feedback of a 0-10 pain rating in person after each stimulation. This procedure was rerun a few times using different starting points and both stimulators. In the second session, 14 trials of randomised painful stimuli were given within the range of lowest perceivable to highest tolerable current level determined in session 1. Subjects rated each stimulus 1s after receiving it, on a 0-10 VAS scale on screen using a keyboard (as practice to the rating procedure used in the task). To determine the final current level to use, a Weibull and Sigmoid function were fitted to session 2’s stimuli and ratings, and current levels for VAS = 1 and 8 were used for low / high pain stimulus for the experiment respectively. The same procedure was repeated for Day 2, and the new fitted current levels were used.

### fMRI data acquisition (Day 1 and 2)

Neuroimaging data was acquired with a 3T Siemens Prisma scanner with the standard 64 channel phased array head coil. Whole-brain functional images were collected with a single echo EPI sequence (repetition time TR=2000ms, echo time TE=26ms, flip angle=80, field of view=240mm), 33 contiguous oblique-axial slices (voxel size 3.2 × 3.2 × 4 mm) parallel to the AC-PC line were acquired. Whole-brain high resolution T1-weighted structural images (dimension 208 × 256 × 256, voxel size 1 × 1 × 1 mm) using standard MPRAGE sequence were also obtained.

### Decoder construction (Day 1)

#### Preprocessing

All preprocessing were conducted using SPM12 (http://www.fil.ion.ucl.ac.uk/spm/software/spm12/) in MATLAB (The MathWorks Inc., Natick, MA, USA).

All functional images were realigned and resliced to the reference functional volume (the first baseline TR after the first 3 dummy TRs obtained in the first session on Day 1). Structural T1 images were coregistered and segmented according to the canonical single subject T1 images. The resulting inverse transformation matrix was used to normalise the bilateral insula ROI obtained from the Automated Anatomical Labeling (AAL) atlas from MNI space to individual subject space. These warped ROI images were then coregistered to the reference functional TR.

#### Feature extraction

Time series were extracted from all voxels within the individual’s insula ROI. To account for BOLD delay and to minimise motion contamination, the times series from TR 3-5 (4-10s) were used from each trial, the first two TRs (0-4s) immediately following pain stimulus were omitted. For denoising, the 5 TRs following 3 dummy TRs at the beginning of each session were used as baseline, each trial ROI time series were normalised by subtracting session baseline mean and divided by baseline standard deviation, then the mean across the TR 3-5 from all trials were extracted for classifier training.

#### Decoder training

Mean insula voxel activity as feature and high/low pain delivered as label were aggregated across all trials within participant for decoder training. Binary classification by Sparse Logistic Regression (SLR, version 1.51) with variational parameters approximation (Yamashita et al., 2008) was used. This results in a sparse matrix of weights for about 5 percent of all voxels within the given ROI. By multiplying weights with feature/voxel intensity signals, the decoder produces the probability of observing current label given trial features (referred as (P(pain) from here, P(pain)=1 means highly likely to have received high pain, P(pain)=0 means unlikely to have received high pain, or highly likely to have received low pain).

For decoder training, all trials were used for training. To estimate decoder accuracy, all trials were partitioned into 10 equal sets with 9 sets for training and 1 set for testing (10 fold cross-validation). The average testing accuracy of 10 iterations of cross-validation were used as estimated decoder accuracy (Tables 1). Trained decoder was tested with another day’s data using the experimental setting.

### Neurofeedback adaptive control algorithm (Day 2)

To allow automated adaptive control of pain stimulus delivery, we used a simple reinforcement learning algorithm (Sutton and Barto, 1998) to update the value of high/low pain states trial-by-trial:

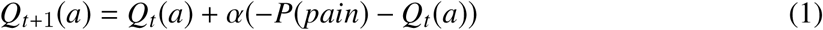

where *t* represent trials, *Q* is the value of given state, *a* is the actions available for the algorithm (i.e. either giving high or low pain, collectively shown as action set *A*), α is learning rate fixed at 0.5.

P(pain) is the decoder-generated probability of current trial’s stimulus being high pain. It’s scaled between [−1,1] when used in the updating function. Higher P(pain) would decrease the value of current pain state more and vice versa, while the value of un-chosen state remained unchanged.

The algorithm selects which pain level to deliver for the next trial using *ϵ*-greedy action selection rule based on current values:

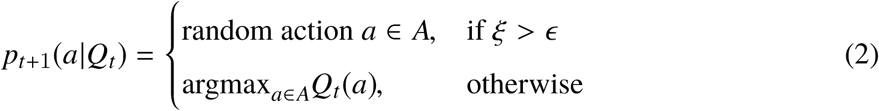

where *ϵ* is the explore ratio fixed at 0.4 (i.e. exploring by choosing a random action by either giving high or low pain 40% of the time, exploiting the other times), *ξ* is a uniform random number drawn within [0,1] at each trial. The random exploration allows a sufficient proportion of alternative pain level to be delivered, to ensure the next participant who uses current participant’s Day 2 sequence to have enough trials of both high and low pain for decoder construction. We also set values to be 0 for both states at the beginning of each session.

### Frequency learning model

The frequency learning model *M* assumes a participant estimates the posterior distribution of a given stimuli *θ* from a previously observed sequence of two possible stimuli *y*_1:*t*_ (i.e. high or low pain) using Bayesian updating (Mars et al., 2008; Meyniel et al., 2016).

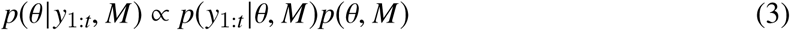

Given the experiment design, participants are assumed to have uninformative prior over the two stimuli at the beginning of each session, which can be represented by a Beta distribution with parameters [1,1]. Since the product of two Beta distributions results in a Beta distribution, the posterior distribution depends only on the frequency of the high and low stimuli *N_h_*, *N_l_*, which has an analytical solution. The posterior mean of the predicted high pain distribution is:

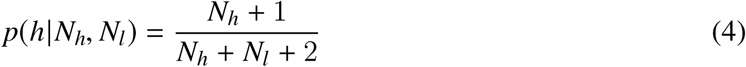

and *P*(*l*|*N_h_*, *N_l_*) = 1 − *p*(*h*|*N_h_*, *N_l_*) given the reciprocal relationship between high/low pain stimuli.

The uncertainty/surprise of current stimulus *h*/*l* at trial *t* can be estimated as the entropy *H* of the posterior mean before updating from trial *t* − 1:

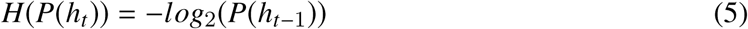

This model does not require model fitting, as participants were assumed to cumulate stimulus counts over the entire session (30 trials), where we assumed perfect memory retention. It is possible to limit the number of trials for frequency memory, or introduce a forgetting ‘leaky factor’ to discount previously experienced trials. However, given that we had no other behavioural data for fitting apart from selective pain ratings, and relatively short sessions, we decided to use the simplest frequency model without fitted parameters.

To determine any learning effects on subjective ratings, we followed the method in Woo et al. (2017)to use subjective rating residuals for correlation analysis with learning model predictors. We regressed subjective ratings with a matrix of high/low pain stimulus identities (high=1, low=-1), and session numbers (1-6) for each individual to obtain rating residuals. The fluctuation of the resulting residuals can be interpreted as modulatory effects on pain beyond the level of nociceptive inputs.

#### Behavioural data

Behavioural data were analysed using Python 3.6, with pandas 0.19.2, scipy 0.18.1, afex 0.16-1.

### fMRI data offline analyses

#### Preprocessing

For offline analysis, functional images were preprocessed using the fmriprep software (build date 21/05/2017, pypi version 0.4.4, freesurfer option turned off, https://github.com/poldracklab/ fmriprep), a pipeline that performs slicetime correction, motion correction, field unwarping, normalisation, field bias correction, and brain extraction using a various set of neuroimaging tools available. The confound files output by fmriprep include the following signals: mean global, mean white matter tissue class, three FSL-DVARS (stdDVARS, non-stdDVARS and voxel-wise stdDVARS), framewise displacement, six FSL-tCompCor, six FSL-aCompCor, and six motion parameters (matrix size 24 × number of volumes). Resulting functional images were smoothed with an 8mm Gaussian kernel in SPM12, except for those in used searchlight analysis.

#### fMRI GLM model

All event-related fMRI data were analysed with GLM models constructed using SPM12, estimated for each participant in the first level. Stick functions at pain stimulation onset were convolved with a canonical hemodynamic response function (HRF). We also included rated trials (duration=10s, from beginning until ITI) as regressor of no interest, in addition to the 24 columns of confound matrix output by fmriprep. Day 1 and 2 data were included in the same GLM, but first-level contrasts were estimated separately for days.

#### Whole-brain comparison (Fig 3)

2 regressors: high/low pain onset (duration=0).

#### Switch trials differences (Fig 5)

4 regressors: trials stimulus different from or identical to that of the previous trial were labelled as switch or non-switch trials, separately for high/low pain (HH, LL, LH, HL), at pain onset (duration=0).

#### Frequency learning posterior probability and entropy (Figs 6b, 6c, 6d)

3 regressors at pain onset (duration=0) with parametric modulators: posterior probability of current stimulus (updated prediction), entropy of previous posterior probability of current stimulus (uncertainty of prediction before updating), actual identity of stimulus (high pain=1, low pain=-1). All parametric modulators mean centred within session, SPM orthogonalisation for these 3 regressors were turned off.

For correction for multiple comparison here and in all analyses, we use whole brain correction or ROI based correction based on a priori hypotheses as appropriate, and the details appear in Table 2. For ROI analyses, we used anatomical binary masks generated using the Harvard-Oxford Atlas (Desikan et al., 2006) (freely available with the FSL software, https://fsl.fmrib.ox.ac.uk/fsl/fslwiki/Atlases), and periaqueductal grey probabilistic atlas (Ezra et al., 2015) for small volume correction. All probability maps were thresholded at 50%, and all masks were applied separately, not combined. We used the frontal medial cortex mask as approximation for VMPFC. Bilateral masks for vlPAG and lPAG were combined respectively. We also used the pgACC peak identified in our previous study of active relief learning (Zhang et al., 2018) for the 8mm spherical ROI mask (sphere peak used: [6,40,12]). We reported all results with p<0.05 (FWE cluster-level corrected, using a p<0.001 cluster-forming threshold (Eklund et al., 2016)), with the exception of searchlight analysis results (MFG/DLPFC SVC had p=0.06, see Table 2).

**Table 2:** Multiple correction (cluster-forming threshold of p <0.001 uncorrected unless stated otherwise, regions from Harvard-Oxford, PAG probabilistic atlas, and previous study. *FWE cluster-level corrected. n=19. H: high pain, L: low pain)

| p* | k | T | Z | MNI coordinates (mm) |  |  | Region mask |
| --- | --- | --- | --- | --- | --- | --- | --- |
|  |  |  |  | x | y | z |  |
| Fig 3: Whole brain comparison (D2>D1, L>H, p<0.005) |  |  |  |  |  |  |  |
| 0.032 | 8 | 4.38 | 3.57 | -16 | -7 | -18 | Amygdala L |
| 0.021 | 28 | 3.81 | 3.22 | -22 | 9 | 6 | Putamen L |
|  |  | 3.62 | 3.10 | -19 | 6 | 2 |  |
|  |  | 3.62 | 3.09 | -26 | -1 | 2 |  |
| 0.040 | 18 | 3.47 | 3 | 23 | 9 | 6 | Putamen R |
|  |  | 3.29 | 2.88 | 26 | 15 | -2 |  |
| Fig 3: Whole brain comparison (D2>D1, H>L, p<0.005) |  |  |  |  |  |  |  |
| 0.034 | 1 | 3.14 | 2.76 | -3 | -30 | -6 | dlPAG (Ezra et al., 2015) |
| 0.034 | 1 | 2.94 | 2.62 | 3 | -30 | -6 |  |
| 0.036 | 1 | 3.14 | 2.76 | -3 | -30 | -6 | lPAG (Ezra et al., 2015) |
| 0.036 | 1 | 2.94 | 2.62 | 3 | -30 | -6 |  |
| Fig 4: Searchlight analysis - decreased information content (D2>D1) |  |  |  |  |  |  |  |
| 0.048 | 2 | 3.94 | 3.3 | -42 | 3 | -2 | Insula L |
| 0.061 | 2 | 4.41 | 3.59 | -38 | 15 | 42 | Middle Frontal Gyrus L |
| 0.078 | 1 | 4.37 | 3.56 | -38 | 35 | 30 |  |
| Fig 4: Searchlight analysis - increased information content (D2>D1, p<0.005) |  |  |  |  |  |  |  |
| 0.045 | 5 | 3.50 | 3.02 | 6 | 44 | 14 | 8mm pgACC sphere at [6,40,12]<br>(Zhang et al., 2018) |
| Fig 5: Whole brain comparison (D2, HL>LL) |  |  |  |  |  |  |  |
| 0.014 | 2 | 4.41 | 3.59 | -26 | -4 | -14 | Amygdala L |
| 0.008 | 6 | 4.81 | 3.81 | 26 | -7 | -14 | Amygdala R |
| Fig 5: Whole brain comparison (D2, LL >HL) |  |  |  |  |  |  |  |
| striatum did not survive SVC |  |  |  |  |  |  |  |
| Fig 6: Frequency learning model - posterior probability of low pain (D2) |  |  |  |  |  |  |  |
| 0.007 | 10 | 4.44 | 3.6 | 0 | 51 | -14 | Frontal Medial Cortex |
| Fig 6: Frequency learning model - entropy (D2) |  |  |  |  |  |  |  |
| 0.039 | 5 | 5.30 | 4.06 | 10 | 41 | 10 | Cingulate Anterior |
| 0.033 | 6 | 4.36 | 3.56 | 0 | 3 | 38 |  |
| 0.002 | 14 | 5.91 | 4.35 | 13 | 41 | 14 | 8mm pgACC sphere at [6,40,12]<br>(Zhang et al., 2018) |
| 0.002 | 31 | 5.24 | 4.03 | -38 | -7 | 2 | Insular cortex (bilateral) |
| 0.032 | 6 | 4.60 | 3.69 | 39 | -4 | 6 |  |

### ROI analysis

Beta estimates were extracted from activation ROIs (see text for mask details). Beta values plotted were the average of all voxels within ROI masks, with statistics showing subject-level SEM. Post-hoc repeated measure ANOVA were conduced with the R package ‘afex’. All t-tests performed were two-tailed. pgACC responses were overlaid on subject-averaged anatomical scans using MRIcroGL (https://www.nitrc.org/projects/mricrogl/). We used voxel-wise correction for multiple comparisons within the ROIs: the insula (required by the task paraigm itself, and the pgACC and PAG given their proposed role in endogenous control (Zhang et al., 2018).)

### Decoder comparison

Decoders were constructed using Day 2 data with the same procedure as Day 1 (Fig 4). This was done to determine whether the decoding performance of insula ROI remained the same, or whether any learning-induced changes might have changed the decoder properties.

Whole-brain searchlight analysis was conducted using the Decoding Toolbox (TDT, v3.98) in MATLAB (Hebart et al., 2015). The toolbox can conduct multivariate decoding analyses at combined trial types within fMRI runs, by extracting features from beta images of relevant regressors in the first level GLM analysis output by SPM. This could lead to higher classification accuracy and lower computation time, comparing to single trial decoding.

A searchlight analysis was carried out within a 10mm radius sphere for the whole brain, with high/low pain categories as unsmoothed beta images from each run for individual participant. TDT produced a decoding accuracy map for each voxel using a leave-one-run-out cross validation scheme, which can be interpreted as the local information content of each voxel (Kriegeskorte et al., 2006). The Day 1 and 2 accuracy maps from each individual were then smoothed with a Gaussian kernel of 4mm, and entered into a standard SPM second level paired t-test as in the GLM analysis above. The resulting T map indicates the changes in decodable information used for pain level decoding across days.

## Appendix Participant instructions

### Day 1 (Decoder construction)

Please rest in the scanner. We are looking at your brain’s response to different levels of pain. You don’t have to do anything.

### Day 2 (Adaptive control)

You don’t need to do anything in this task. The computer is trying to work out if you feel pain or not, by looking at your brain activity. If it thinks you felt pain, it will try and change the pain stimulation to stop you from having pain. If it thinks you did not feel much pain, it will try not to change anything. However, it cannot do this very reliably, as reading the brain activity is difficult, so it may often make mistakes.

During your first scan, we gave a random sequence of pain stimuli - some high, and some low. Using this data, we have trained a computer program to tell how much pain you were feeling during each shock, based on your brain activity. It is good, but not perfect - it gets it right about 80% of the time.

In today’s scan, the computer program can influence the pain level you get. If it thinks you felt a lot of pain, it will influence the pain machine to give you less pain in the future. If it thinks you did not feel much pain, it will try to influence the pain machine to continue to give you little pain. In other words, it is trying to help you get less pain! This is a difficult job for the computer program, because it is not perfect at reading your brain activity as soon as it is active (i.e. within a few seconds).

It is up to you what you do in the task. You can do nothing, and hope that the system works well, and the computer learns to reduce the pain. Or you can try to influence the computer using your thoughts, in any way that you like.

### Post-training survey (Day 2)

1. Do you think the machine was successful in reading your pain and trying to reduce it?
2. Did you try to influence the computer by doing or thinking anything?
3. If so, what did you do/think?
4. And if so, do you think you were successfully able to influence it?
5. Any other comments or feedback?

## Acknowledgements

Research was supported by the ‘Application of DecNef for development of diagnostic and cure system for mental disorders and construction of clinical application bases’ of the Strategic Research Program for Brain Sciences from Japan Agency for Medical Research and development (AMED), the Wellcome Trust (UK), Arthritis Research UK, the National Institute for Information and Communications Technology (NICT, Japan), the Japanese Society for the Promotion of Science (JSPS), and S.Z. is supported by the W.D. Armstrong Fund and the Cambridge Trust. K.S. was supported by JSPS Kakenhi Grant Number 17H04789. We thank the imaging teams at the Advanced Telecommunications Research Institute for their assistance in performing the study.

